# The genome sequence of the Montseny horsehair worm, *Gordionus montsenyensis* sp. nov., a key resource to investigate Ecdysozoa evolution

**DOI:** 10.1101/2023.06.26.546503

**Authors:** Klara Eleftheriadi, Nadège Guiglielmoni, Judit Salces-Ortiz, Carlos Vargas-Chavez, Gemma I. Martínez-Redondo, Marta Gut, Jean-François Flot, Andreas Schmidt-Rhaesa, Rosa Fernández

## Abstract

Nematomorpha, also known as Gordiacea or Gordian worms, are a phylum of parasitic organisms that belong to the Ecdysozoa, a clade of invertebrate animals characterized by molting. They are one of the less scientifically studied animal phyla, and many aspects of their biology and evolution are still unknown, partially due to the lack of genomic resources for this phylum. As part of the European Reference Genome Atlas pilot effort to generate reference genomes for European biodiversity, we present the taxonomic description and chromosome-level genome assembly of a newly described species of Nematomorpha (*Gordionus montsenyensis* Schmidt-Rhaesa & Fernández sp. nov.). The final assembly has a total length of 288 Mb in 396 scaffolds with an N50 of 64.4 Mb, 97% of which is scaffolded into 5 pseudochromosomes. The circular mitochondrial genome was also assembled into a 15-kilobases sequence. Gene annotation predicted 10,320 protein-coding genes in the nuclear genome. In this study, we contribute a key genomic resource to not only explore the evolution of Ecdysozoa, but also to further our understanding on the genomic basis of parasitic lifestyles. In addition, we describe a species new to science from this enigmatic animal phyla.

## Introduction

Nematomorpha, also known as Gordiacea or Gordian worms, are a phylum of parasitic organisms that belong to the Ecdysozoa, a clade of invertebrate animals characterized by molting (Aguinaldo et al. 1997). The name “Gordian” is derived from the legendary Gordian knot, as nematomorphs often intertwine themselves into compact balls that resemble knots. These animals can measure up to 1 m in length, with a diameter ranging from 1 to 3 millimeters. There are approximately 360 described species of horsehair worms, but as this is one of the most understudied animal phyla, their true diversity is probably much larger in terms of number of species (Schmidt-Rhaesa 2013). Two classes exist within the phylum, one marine (Nectonematida) and the other freshwater (Gordiida) (Schmidt-Rhaesa 2013). Horsehair worms are commonly found in moist environments such as watering troughs, swimming pools, streams or puddles. While the adult worms are free living either in freshwater or marine environments, the larvae are parasitic and rely on arthropods including beetles, cockroaches, mantises, orthopterans, and crustaceans. The host must be in contact with water for the mature adult to emerge from the body cavity (Hanelt and Janovy 2003). The parasite may alter the behavior of the host and increase the chance that it ends up in water, where the adult exits the body of the host (Thomas et al. 2002). As expected, given their parasitic lifestyle (Hanelt, Thomas and Schmidt-Rhaesa 2005), nematomorphs are characterized by a series of morphological peculiarities such as the loss of circulatory, excretory and digestive systems (for instance, adults have lost their mouth and do not feed - they just reproduce). The sex of an individual and some characters can be mostly recognized with simple optics, but specific determination requires scanning electron microscopy imaging. Structures important for identification are the fine cuticle structure and the cuticular structures on the posterior end of males (Hanelt, Thomas and Schmidt-Rhaesa 2005).

The phylogenetic position of Nematomorpha within the Metazoa Tree of Life has been recalcitrant to resolution (Laumer et al. 2019), hampered not only by the lack of high-quality genomic data for this phylum but also for some of the other phyla composing Ecdysozoa, such as Kinorrhyncha, Loricifera or Priapulida. However, in the majority of cases molecular data have only been used to investigate systematic relationships between or within phyla closely related to Nematomorpha, with hairworm sequence data often solely serving as an outgroup in phylogenetic analyses (Aguinaldo et al. 1997; Blaxter et al. 1998; Sørensen et al. 2008). In this study, we contribute a key genomic resource to unveil not only the evolution of Ecdysozoa, but also to further our understanding on the genomic basis of parasitic lifestyles. In addition, we describe a species new to science from this enigmatic animal phylum.

## 1. Species Description

### Taxonomic Results

Phylum Nematomorpha Vejdovsky, 1886

Order Gordioidea Rauther, 1930

Family Chordodidae, 1900

Genus *Gordionus* Müller, 1926

*Gordionus montsenyensis* Schmidt-Rhaesa & Fernández sp. nov.

### Material examined

Holotype. Adult male, anterior and posterior end (accession number ZMH V13667 in the Zoological Museum Hamburg (ZMH) of the Leibniz Institute for the Analysis of Biodiversity Change (LIB), Hamburg, Germany), collected from the Tordera river on the foothill of Montseny Natural Reserve, Barcelona (Spain)(code Nem3 in Fig. 2). Paratype. One female, full body (accession number ZMH V13668), with the same collection data of the holotype. Additional material: two female adults, full body, one female adult, anterior and posterior end (accession numbers ZMH V 13669 to 13671)(codes Nem4 and Nem6 in Fig. 6). DNA extracted and kept at the cryocollection of the Metazoa Phylogenomics Lab, Institute of Evolutionary Biology (Spanish National Research Council - University Pompeu Fabra).

### Molecular data

Genbank accession numbers: COI - OR136460-OR136462 (NCBI).

Specimens varied from 10 to 17 cm in length (Fig. 1A). All specimens were completely white, with lack of pigmentation. The cuticle of both sexes is structured into polygonal structures called areoles (Fig. 1B,C). Areoles are separated by a groove, no further structures (such as bristles) were observed within these grooves. The surface of the areoles is smooth. The posterior end of the male has two tail lobes, the cloacal opening is ventrally, about 50 µm anterior of the start of the bifurcating tail lobes (Fig. 1B,D). The entire posterior end is covered by areolas with the exception of the ventral side (Fig. 1B). On the ventral side of the tail lobes and around the cloacal opening, the surface is smooth or areolas are only indicated as shallow elevations without grooves between them. More anteriorly, in the region of the adhesive warts (Fig. 1B,E), areoles are more or less rectangular and have fine transverse grooves.

**Figure 1.**
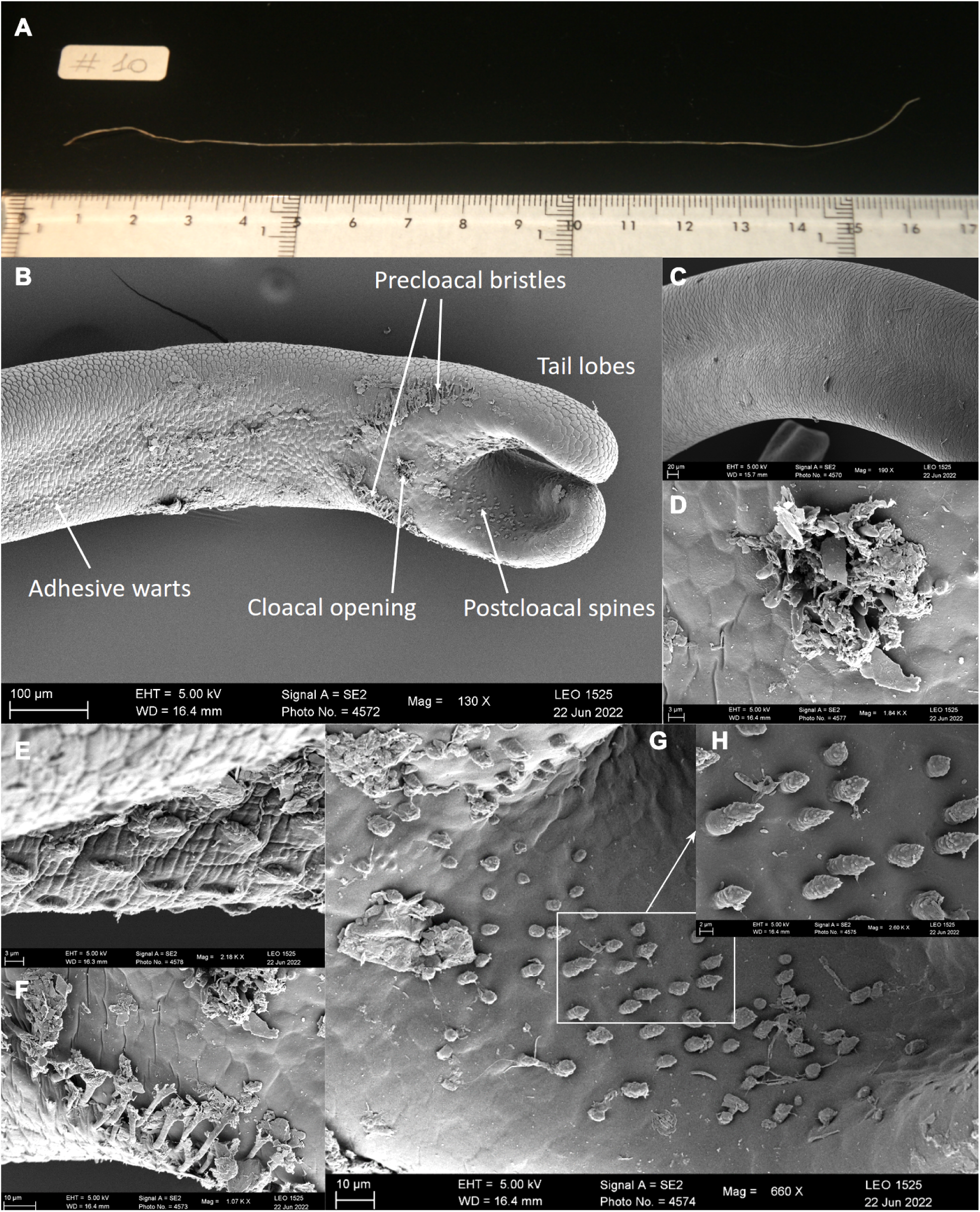
*Gordionus montsenyensis* sp. nov. 1A. Picture of a fresh specimen. 1B-H. Electron microscopy images showing the posterior end of a male specimen (holotype). 1B. Posterior end with main morphological features indicated. 1C. Cuticle. 1D. Cloacal opening. 1E. Adhesive warts. 1F. Precloacal bristles. 1G. Postcloacal spines. 1H. Detail of the postcloacal spines.

The male posterior end includes cuticular structures in the form of circumcloacal spines, precloacal bristles, postcloacal spines and adhesive warts (Fig. 1B). Twelve to maximally 15 stout spines surround the cloacal opening (their exact number could not be counted because some are covered by dirt or sperm) (Fig. 1D). Their apical end is rounded and not pointed. Anterolaterally of the cloacal opening is a pair of oblique rows of bristles, which are directed posteriorly (Fig. 1F). Apically, these bristles branch into several “arms”. On the inner side of the tail lobes are postcloacal spines. They are pointed and have an irregular, “warty” surface. At least in some spines the pointed tip consists of a tube-like structure (Fig. 1G,H).

Phylogenetic analysis recovered *G. montsenyensis* as a clade within the genus *Gordionus* in sister-group position to *Gordionus alpestris* with strong support (Fig. 2). In addition, the mitogenome from *G. montsenyensis* resulted to be highly similar to that of *G. alpestris* (Fig. 3), and their pairwise similarity was 95.04% (Fig. 3). These results support the assignment of this new species to the genus *Gordionus,* supported as well by morphology.

**Figure 2.**
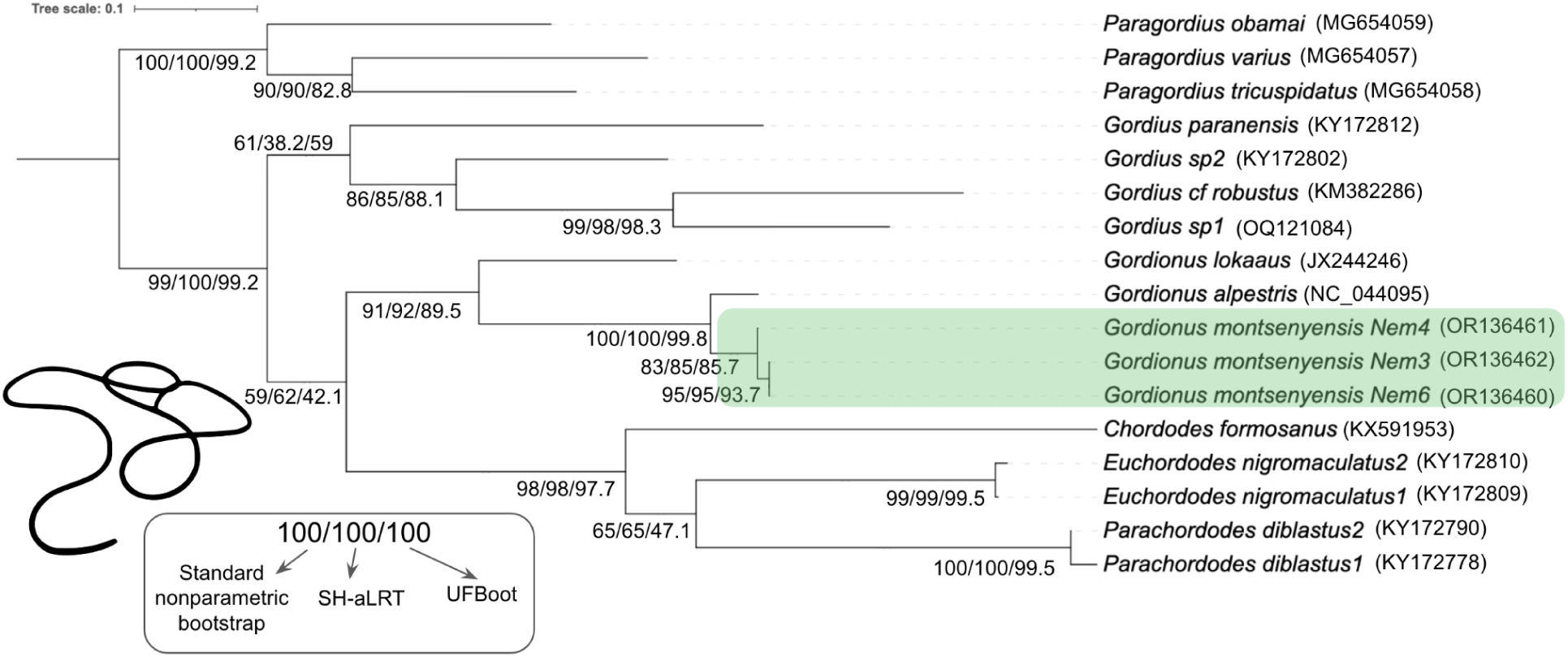
Maximum likelihood phylogenetic tree showing the placement of *Gordionus montsenyensis* sp. nov. Standard nonparametric bootstrap, SH-aLRT and UFBoot support is indicated for each node. NCBI accession numbers are indicated in brackets. Nematomorph silhouette retrieved from PhyloPic (credit: Eduard Solà Vázquez, vectorised by Yan Wong).

**Figure 3.**
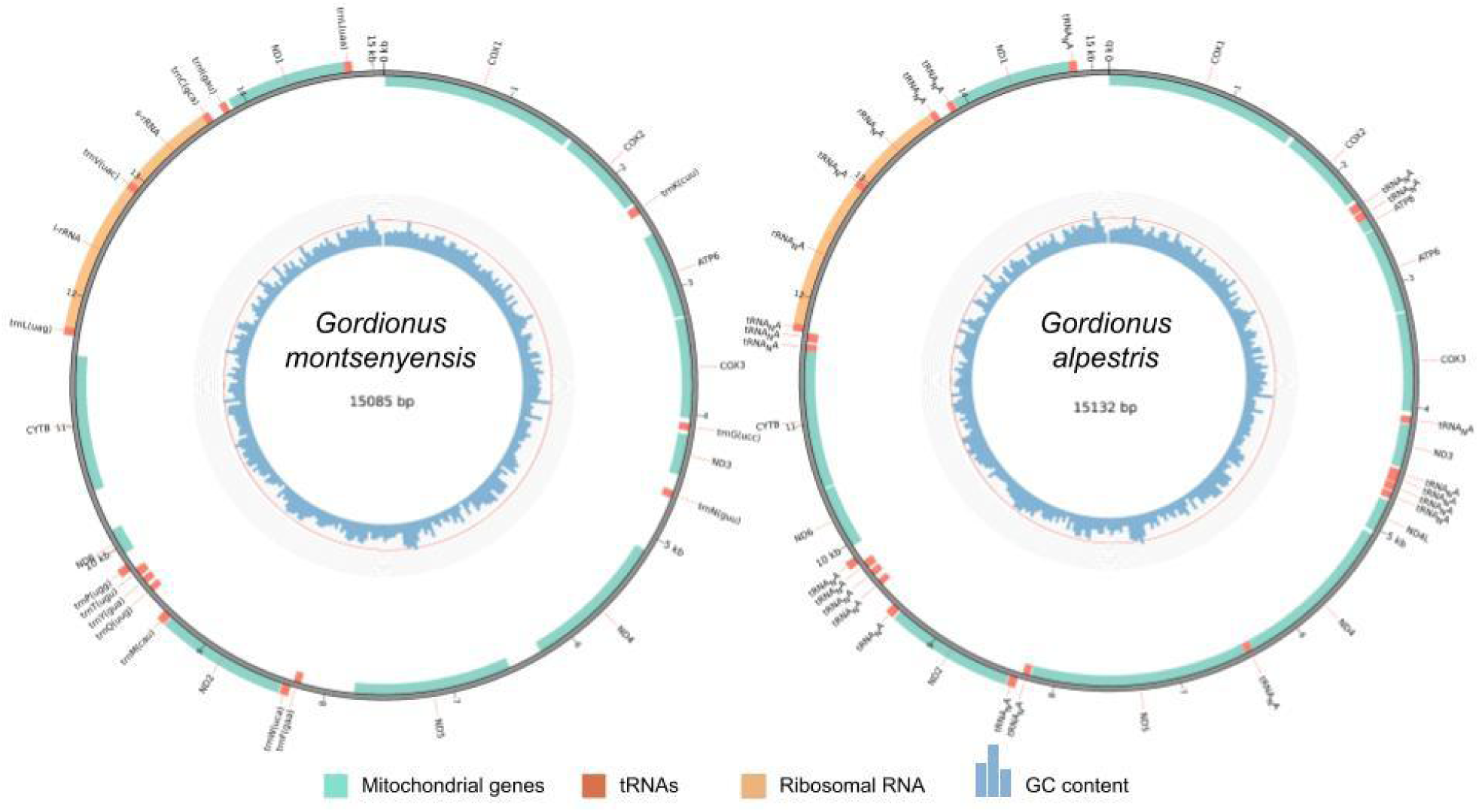
Mitogenome annotation of *Gordionus montsenyensis* (left) and *Gordionus alpestris* (right). Position of mitochondrial genes, tRNAs, ribosomal RNA and GC content are indicated.

## Discussion

Our understanding of the nematomorph fauna in Spain is limited and fragmented.. It includes seven species (de Villalobos et al. 2001, Schmidt-Rhaesa and Cieslak 2008, Schmidt-Rhaesa and Martínez 2016): *Gordionus barbatus* Schmidt-Rhaesa and Cieslak, 2008, *Gordionus wolterstorffii* (Camerano, 1888), *Gordius aquaticus* Linné, 1758, *Gordius gonzalezi* Schmidt-Rhaesa and Martínez, 2016, *Gordius plicatulus* Heinze, 1937, *Paragordionus ibericus* Schmidt-Rhaesa and Cieslak, 2008, *Paragordius tricuspidatus* (Dufour, 1828). Four additional species were listed by Gerlach (1978): *Gordionus violaceus* (Baird, 1853), *Parachordodes gemmatus* (Villot, 1885), *Parachordodes speciosus* (Janda, 1894) and *Parachordodes tolosanus* (Dujardin, 1842). These four records are likely misinterpretations. They all correspond to having been collected in Galicia, without any more specified location. It is likely that the Central European region (today partly in Poland and Ukraine) is meant and not the Spanish province.

In general, the morphology of the specimens correspond well to the characters of the genus *Gordionus* (see, e.g. Schmidt-Rhaesa 2013): one type of areoles is present and in the male posterior end there are precloacal bristles, postcloacal spines, circumcloacal spines and adhesive warts. This result was corroborated with the molecular biology analysis, as *G. montsenyensis* clustered together with other species from the same genus (Fig. 2) and the mitogenome was very similar to that of *G. alpestris* (pairwise identity between mitogenomes is 95.04%; Fig. 3). Most *Gordionus* species have fine bristles in the interareolar spaces, but a few species including *G. montsenyensis* lack such bristles: *G. bageli* (Schmidt-Rhaesa and Gusich 2010), *G. lineatus* (Smith 1991, Schmidt-Rhaesa et al. 2003, 2009), *G. lokaaus* (Begay et al. 2012) and probably *G. diligens* (de Villalobos et al. 1999). Other characters (furcated precloacal bristles, stout circumcloacal spines and elongate adhesive warts) are similar to those in other *Gordionus* species, but the shape of the postcloacal spines in *G. montsenyensis* is unique. In all other species documented by SEM, the spines are pointed and have a smooth surface (e.g. Schmidt-Rhaesa and Cieslak 2008 for *G. turkensis*, Begay et al. 2012 for *G. lokaaus*), therefore the warty appearance of the spines in *G. montsenyensis* is unique and justifies the description as a new species.

Nematomorphs are not very rich in characters that can be used for identification. Besides the cuticular structure, the posterior end of the males is the most important body part for identification. The female specimen was confirmed to belong to *G. montsenyensis* through DNA barcoding, the similar cuticular structure, size and color and the same sampling location, but the female alone does not have any species-specific characters. Horsehair worms are often collected in low numbers and many species were described on the basis of few specimens. Although it is desirable to have more specimens and document some intraspecific variability of characters, often only single or few specimens are available.

## 2. Genome sequence report

A total of 340-fold coverage of Oxford Nanopore long reads with a read N50 of 18,184 basepairs (bp) and 80-fold coverage in 154.9 millions paired 151-bp DNA Illumina reads were generated. The coverage was estimated against the size of the final genome assembly generated. A Hi-C library of 102.9 millions paired reads was constructed for further scaffolding into chromosomes the draft genome assembly. The species appears to be diploid, with genome length of 279 Megabases (Mb) and 0.548% of heterozygosity as predicted in the *k*-mer distribution analysis (Fig. 4). Manual curation of the assembly corrected 8 misjoins.The final assembly has a total length of ∼288 Mb in 396 scaffolds with a scaffold N50 of 64.4 Mb (Table 2), a haploidy score of 0.985 indicating proper haplotype collapsing further supported by *k*-mer analysis, and 97.05% of the assembly sequence assigned to 5 pseudochromosomes (Figs. 5-9). The BlobPlot showing the GC coverage of the final manually curated assembly against the NCBI-nt showed potential contaminant sequences from Arthropoda, Mollusca or Annelida. None of these sequences were filtered, since due to the parasitic life cycle of nematomorphs, these sequences could be horizontally transferred sequences. Also, these sequences could be artifacts as a consequence of the underrepresentation of the phylum to the database (Figs. 7-8). The GC content spans 26.93% of the total assembly length. The evaluation using BUSCO v5.4.7 (Manni et al. 2021) resulted in completeness of 71.6% against the metazoa_odb10 reference set (Fig. 9). The mitochondrial genome spans 15,085 bp and it has been deposited under a different accession number (Fig. 3). Up to date there are two more genomes of the phylum of Nematomorpha (Cunha et al., 2023), one of them chromosome-level. A comparison of the 3 publicly available to the scientific community genomes can be shown in Table 1.

**Figure 4.**
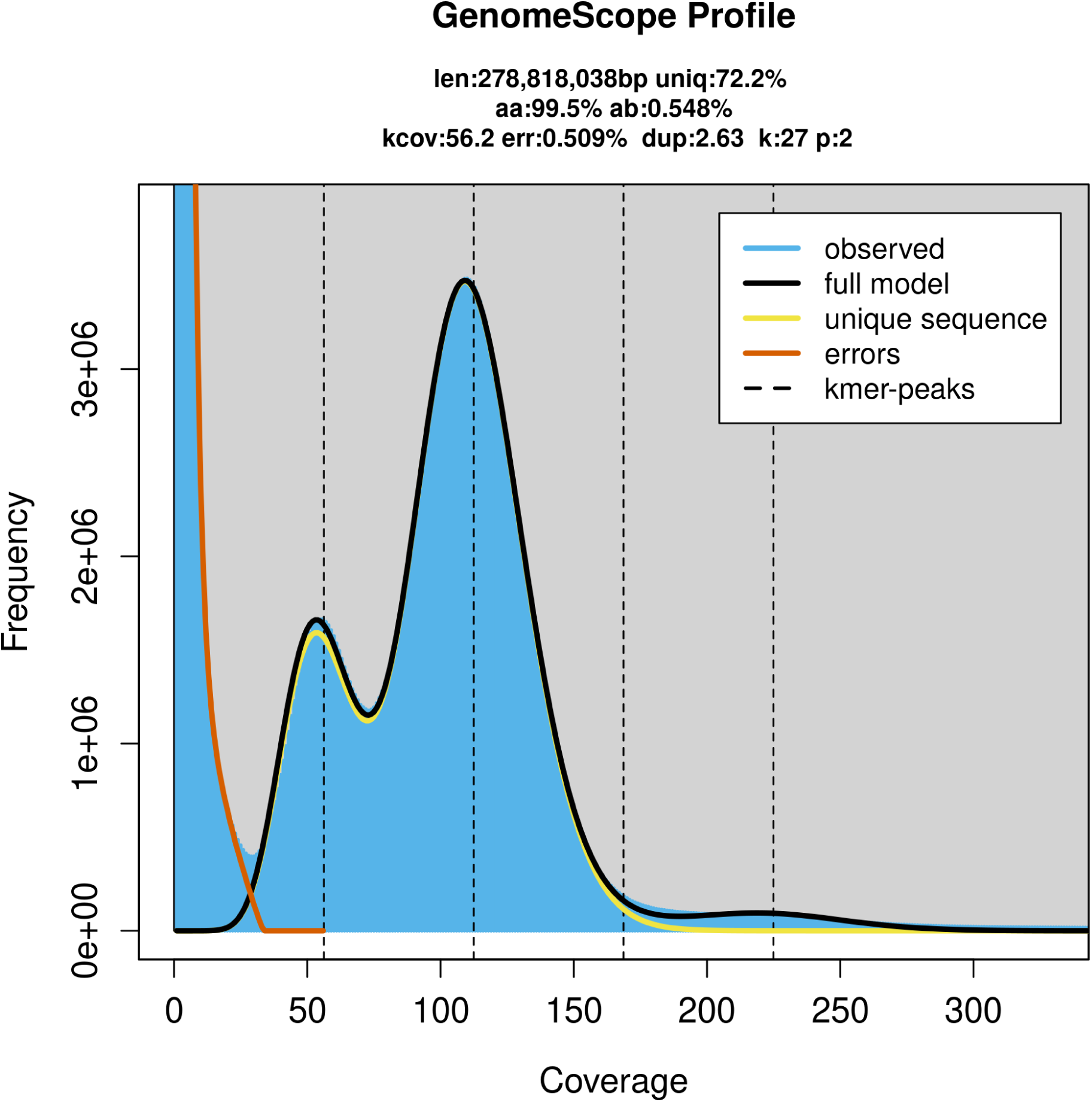
Genome profiling showing the estimated genome size, the heterozygosity and the ploidy of the genome of *Gordionus montsenyensis* sp. nov. based on *k*-mer distribution using GenomeScope2. The y-axis represents the k-mers frequency and the x-axis the k-mer coverage. The peaks indicate a diploid species.

**Figure 5.**
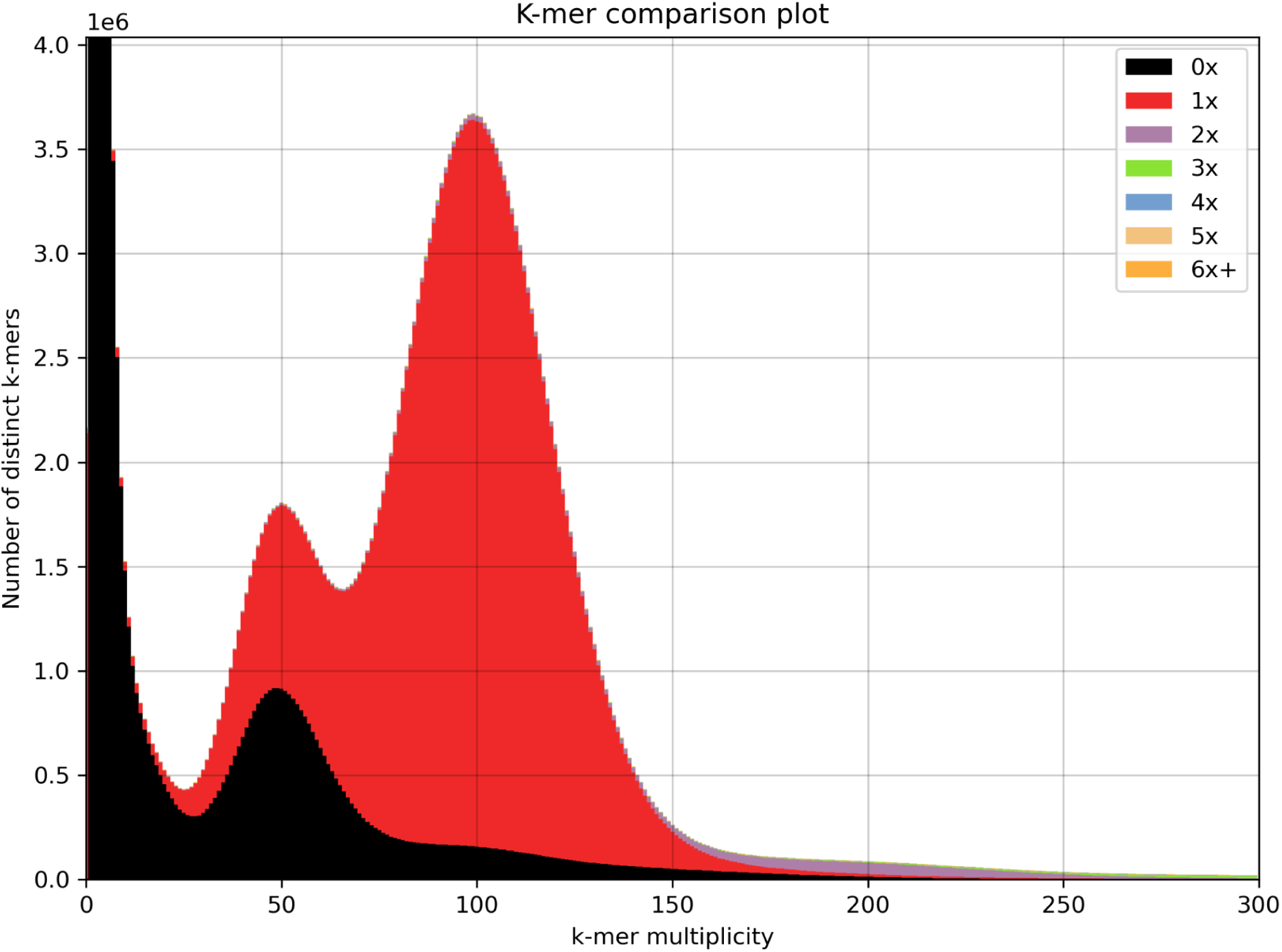
*k*-mer analysis of the Illumina DNA reads against the final assembly using KAT.

**Figure 6.**
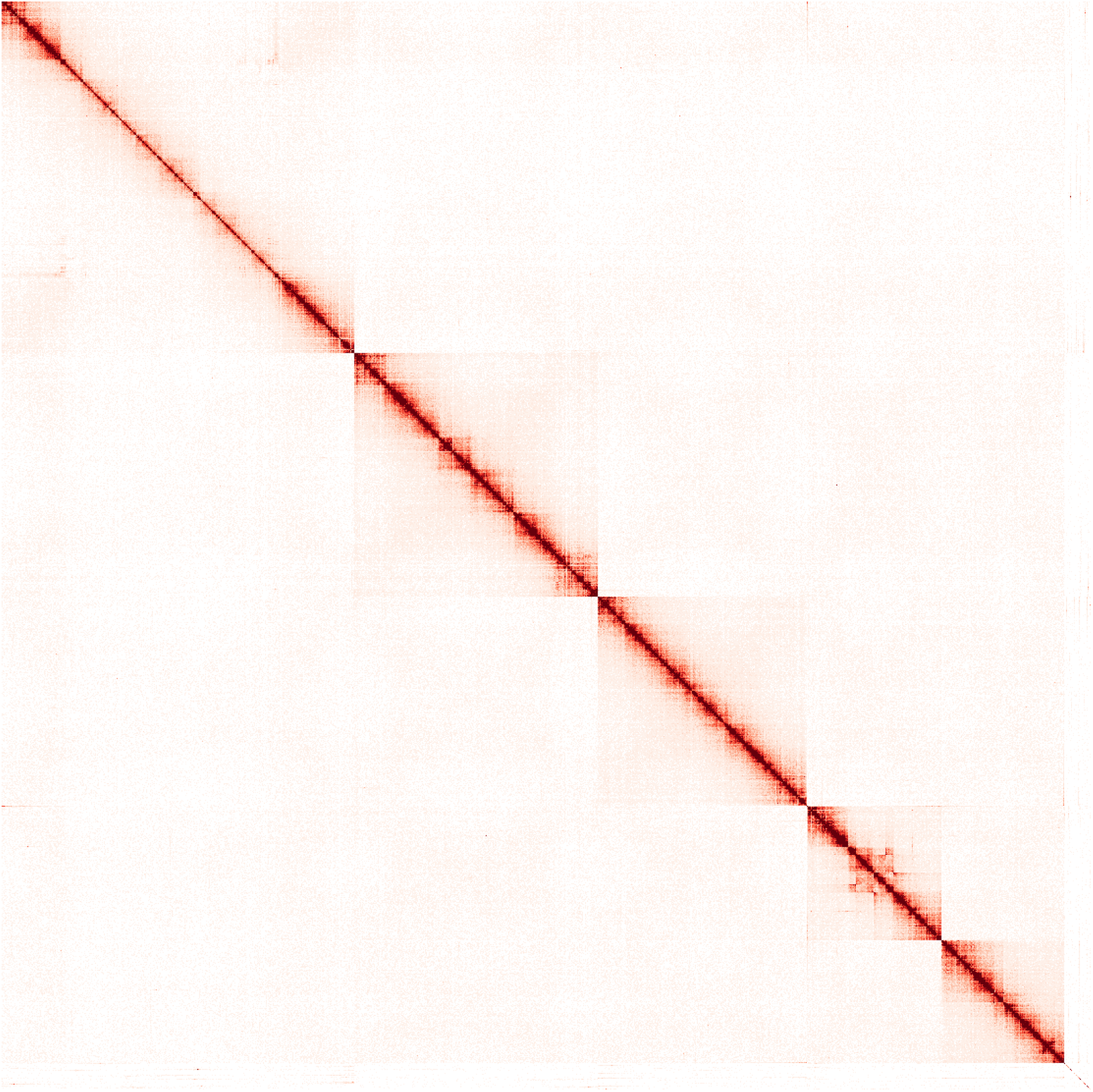
Hi-C contact map of the final manually curated genome assembly of *Gordionus montsenyensis* sp. nov. visualized using the hicstuff pipeline.

**Figure 7.**
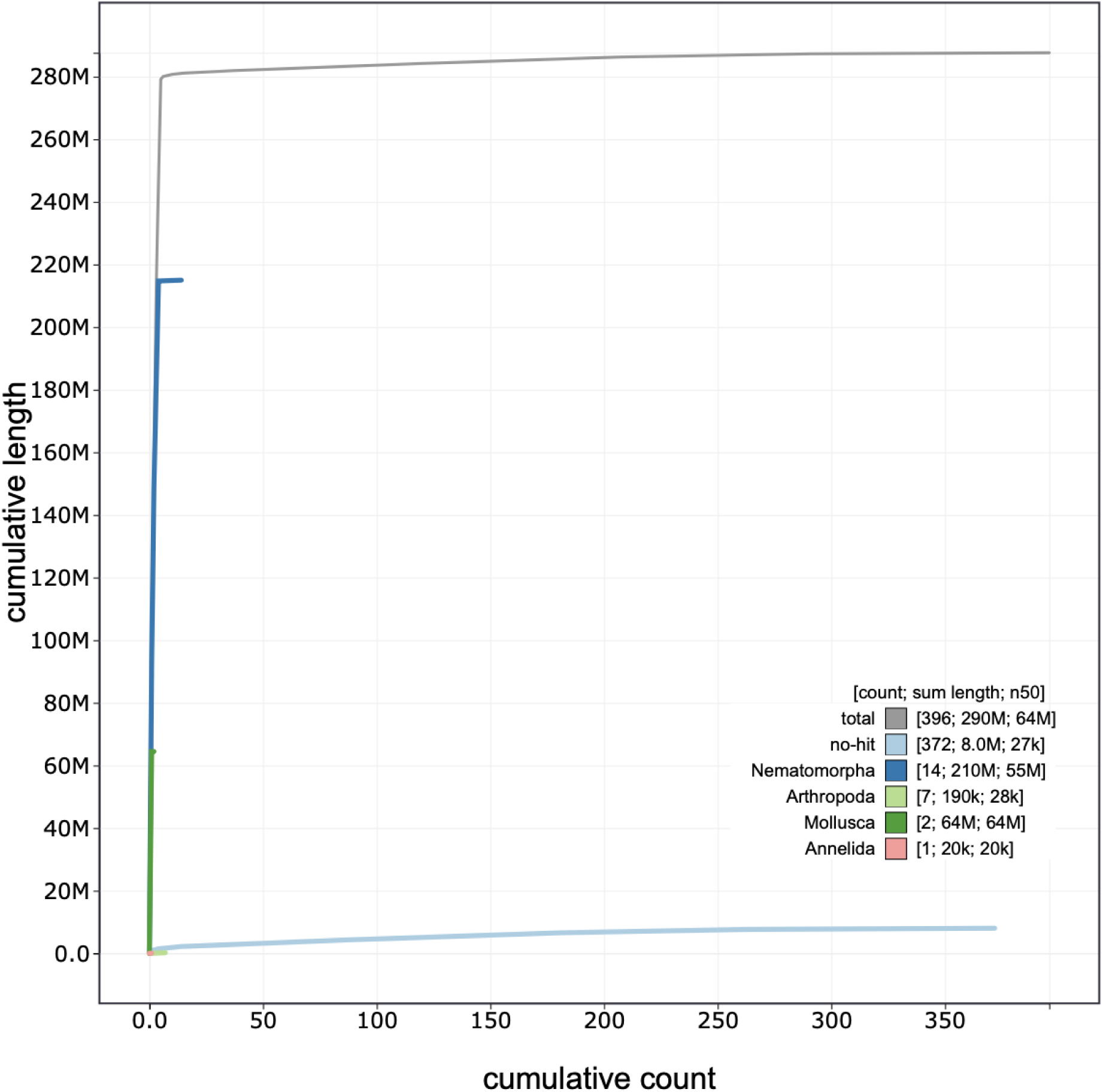
BlobToolKit cumulative sequence plot. The grey line represents the cumulative length for the total number of scaffolds. The colored lines represent the cumulative length of each taxonomic assignment using the bestsum taxrule by a blast search against the nt database.

**Figure 8.**
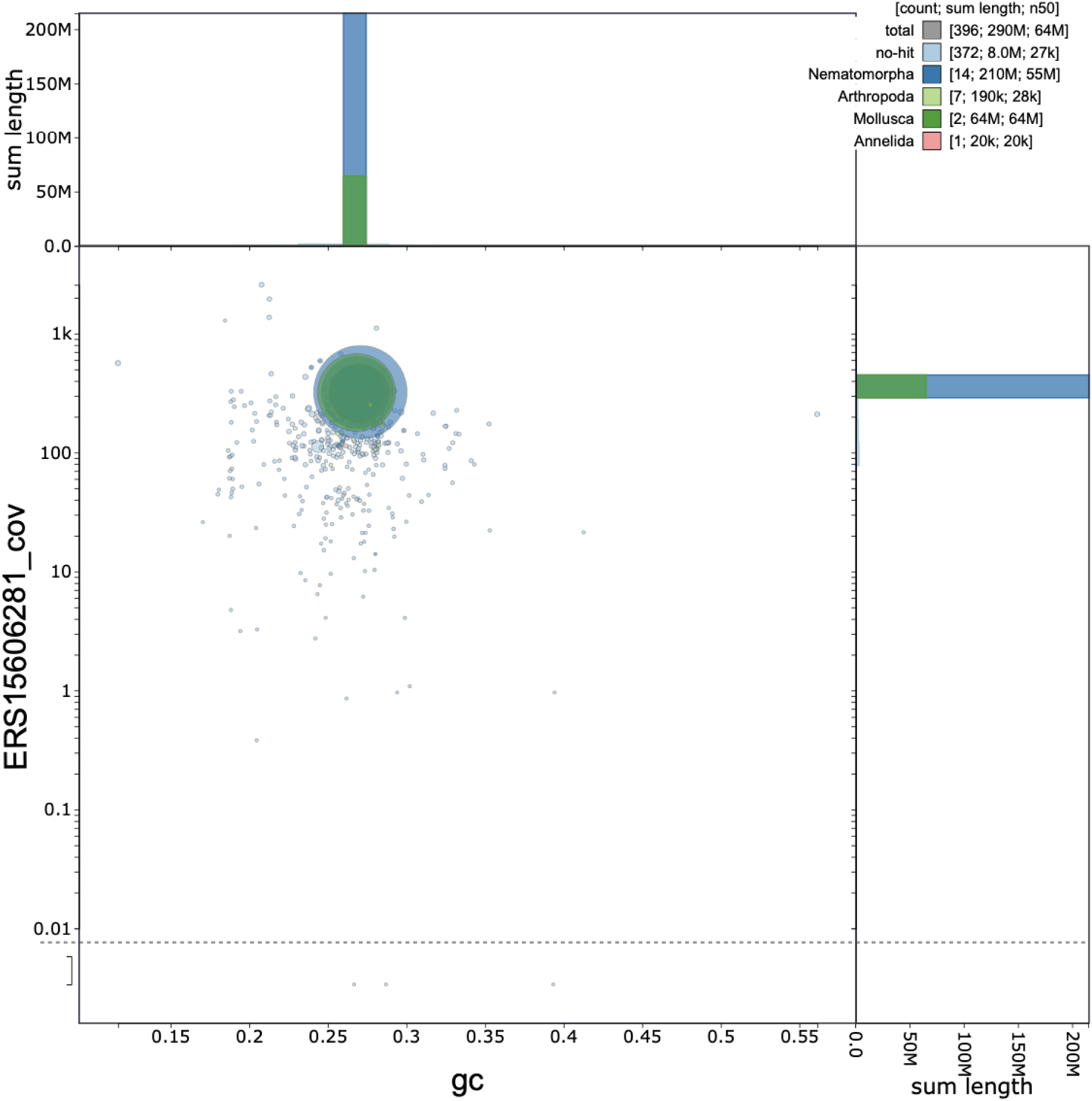
BlobToolKit GC-coverage plot of the final manually curated genome assembly of *Gordionus montsenyensis* sp. nov. The size of each circle is proportional to the scaffold length and each color represents taxonomic assignment by a blast search against the nt database. It should be noted that the *Gordionus montsenyensis* sp. nov. genome is the first genome assembled of the phylum of Nematomorpha, explaining the misidentification of other phyla.

**Figure 9.**
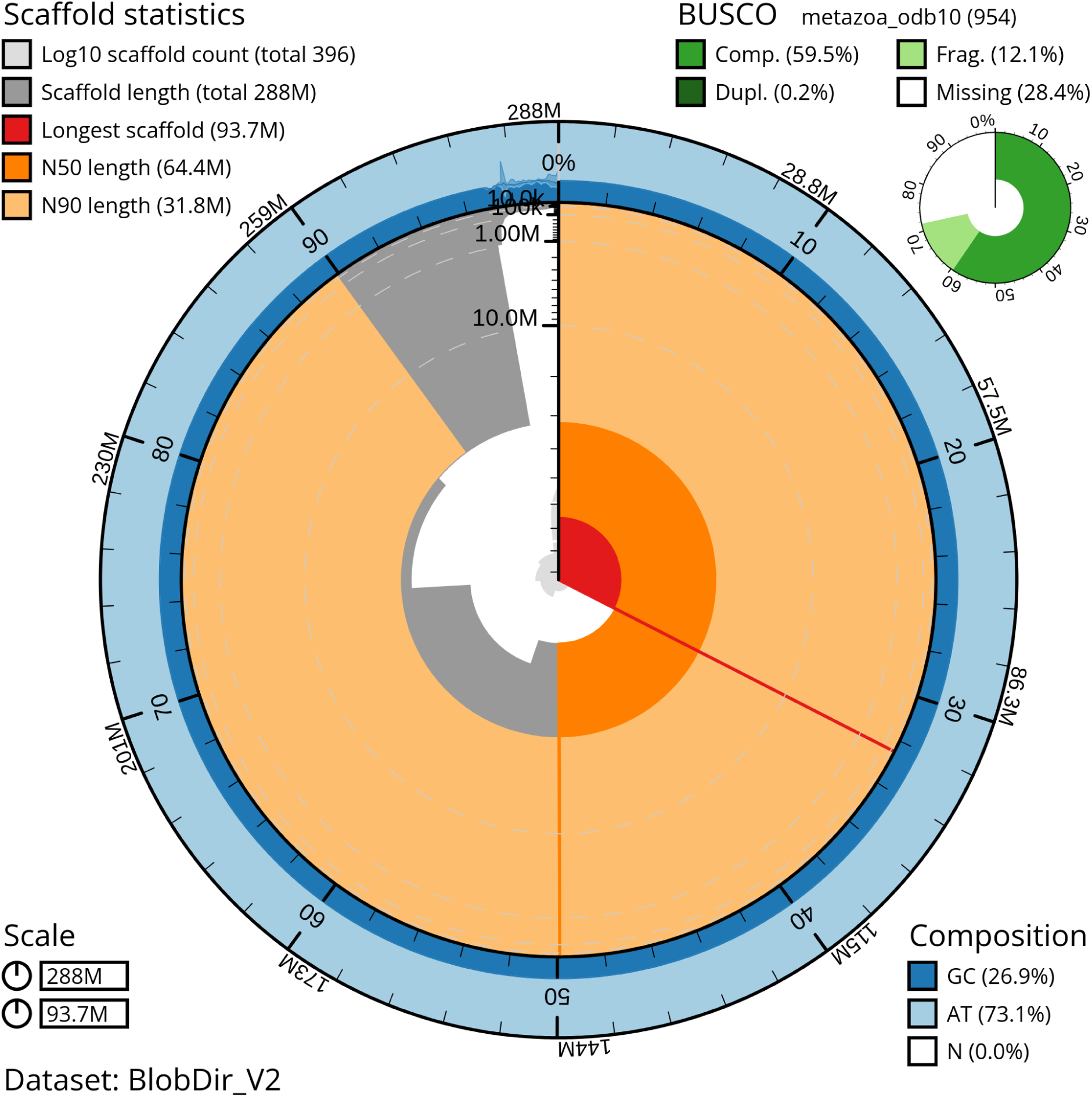
BlobToolKit Snail plot shows summary statistics of the chromosome-level *Gordionus montsenyensis* sp. nov. genome assembly and BUSCO completeness score. The main plot is divided into 1,000 size-ordered bins around the circumference with each bin representing 0.1% of the 287,640,327 bp assembly. The distribution of sequence lengths is shown in dark grey with the plot radius scaled to the longest sequence present in the assembly (93,749,283 bp, shown in red). Orange and pale-orange arcs show the N50 and N90 sequence lengths (64,442,710 and 31,754,433 bp), respectively. The pale grey spiral shows the cumulative sequence count on a log scale with white scale lines showing successive orders of magnitude. The blue and pale-blue area around the outside of the plot shows the distribution of GC, AT and N percentages in the same bins as the inner plot. A summary of complete, fragmented, duplicated and missing BUSCO genes in the metazoa_odb10 set is shown in the top right.

**Table 1.** Comparison of the publicly available genomes of the phylum Nematomorpha. Dashes indicate information not provided in Cunha et al., 2023.

| Metrics | *Gordionus montsenyensis* sp. nov.<br>(This study) | *Acutogordius australiensis*<br>(Cunha et al., 2023) | *Nectonema munidae*<br>(Cunha et al., 2023) |
| --- | --- | --- | --- |
| Total bases | 288 Mbp | 201 Mbp | 213 Mbp |
| GC content | 26.93% | 28.50% | 30.30% |
| Scaffolds number | 396 | 1,323 | 1,061 |
| Longest scaffold | 93 Mbp | 46.6 Mbp | 3.2 Mbp |
| Scaffold N50 | 64.4Mbp | 38 Mbp | 0.71 Mbp |
| Scaffold L50 | 2 | - | - |
| % of genome anchored in X chromosome-scale scaffolds | 97.05% in 5 scaffolds | 83.4% in 4 scaffolds | - |
| Repetitive elements | 63.79% | 49.41% | 53.47% |
| Predicted genes | 10,320 | 11,114 | 8,717 |
| BUSCO (Metazoa odb10) | C:59.5% [S: 59.3%, D: 0.2%], F: 12.1%, M: 28.4% | C:56.2% [S:50.7%, D:5.5%], F:11.5%, M:32.3% | C:59.0% [S:58.4%, D:0.6%], F:9.7%, M:31.3% |

**Table 2.** Project accession data for *Gordionus montsenyensis* sp. nov.

| Assembly identifiers |  |
| --- | --- |
| Species | *Gordionus montsenyensis* |
| Specimen | Nem9 (ONT), Nem7 (Illumina), Nem5 (Hi-C), Nem11 (RNAseq) |
| NCBI taxonomy ID | 3053472 |
| Isolate information | Whole specimen |
| toIID | tfGorSpeb1 |
| *Raw data accessions* |  |
| Oxford Nanopore Technologies | ERS15606281 |
| Illumina | ERS15606282 |
| Hi-C Illumina | ERS15606283 |
| RNA-seq | ERS15606284 |
| *Genome assembly* |  |
| Assembly accession number | PRJEB63266 |
| Mitogenome accession number | PRJEB63265 |
| Span (Mb) | 287.6 |
| Number of scaffolds | 396 |
| Scaffold N50 length (Mb) | 64.4 |
| Scaffold L50 | 2 |
| Scaffold N90 length (Mb) | 31.8 |
| Scaffold L90 | 5 |
| Longest scaffold (Mb) | 93.7 |
| Number of contigs | 571 |
| Contig N50 length (Mb) | 13.4 |
| Contig L50 | 12 |
| Contig N90 length (Mb) | 2.3 |
| Contig L90 | 74 |
| BUSCO* completeness | C:59.5%[S:59.3%,D:0.2%],F:12.1%,M:28.4:%,n:954 |

| *Genome annotation* |  |
| --- | --- |
| Genome annotation file accession number | PRJEB63266 |
| Number of protein-coding genes | 10,320 |
| Number of transcripts | 15,442 |
| *BUSCO scores based on the metazoa odb10 BUSCO set using v5.4.7. C= complete [S= single copy, D=duplicated], F=fragmented, M=missing, n=number of orthologues in comparison. |  |

## 3. Genome annotation report

Repeat annotation pipeline masked 63.79% of the genome, with the highest percentage of repeats belonging to DNA transposons class (Table 3). It is worth noticing the lack of SINE or Penelope repetitive elements. Structural annotation yielded 10,320 protein coding genes and 15,442 transcripts (Table 2).

**Table 3.** Repetitive elements in the genome of *Gordionus montsenyensis* sp. nov.

|  | **Number of elements** | **Length occupied** | **% of sequence** |
| --- | --- | --- | --- |
| *SINEs* | 0 | 0 | 0.00% |
| *Penelope* | 0 | 0 | 0.00% |
| *LINEs* | 28,572 | 14,955,707 bp | 5.20 % |
| *LTR elements* | 38,406 | 23,012,610 bp | 8.00 % |
| *DNA transposons* | 156,757 | 57,712,924 bp | 20.06% |
| *Rolling-circles* | 2,235 | 962,498 bp | 0.33% |
| *Unclassified* | 383,874 | 86,725,537 bp | 30.15% |

## Materials and Methods

### Species collection, electron microscopy and phylogenetic analysis

Twelve specimens were hand-picked in the Tordera river in Montseny Natural Reserve in June 2021. Specimens were half buried in the riverbed or coiled around plant stalks. Six of them (one male and five females) were fixed in 70% EtOH for morphological investigation. The other six were flash frozen for genome sequencing (sex was not recorded).

For scanning electron microscopy (SEM), a piece of 0.5-1mm length was removed from the midbody and in the male the posterior end was cut approximately 1mm from the posterior tip. Pieces were dehydrated in an increasing ethanol series, critically point dried and coated with gold in a sputter coater. Investigation took place with a LEO SEM 1524 and digital images were taken.

For molecular biology investigation, DNA from three individuals (one adult male and two adult females) was extracted from a 1 cm-piece of the central body with the Qiagen Dneasy Blood & Tissue Kit following manufacturer’s instructions. PCR amplification conditions and primers were as in Folmer et al. (1994). DNA amplicons were cleaned using the Clean-Easy PCR & Gel Extraction Kit (Canvax) and sent to Macrogen Inc. for sequencing.

The phylogenetic position of the species was investigated through maximum likelihood inference. For that, we retrieved nucleotide COI nematomorph sequences from NCBI, aligned them with MAFFT with the progressive method L-INS-1 (Katoh and Standley 2013), and inferred a maximum-likelihood tree with IQ-TREE2 (Minh et al. 2020), with automated model selection (Kalyaanamoorthy et al. 2017) (Best-fit model: K3Pu+F+I+G4 chosen according to Bayesian Information Criteria). *Paragordious obamai, varius* and *tricuspidatus* were used as outgroups. Three analyses were run with different branch support methods: (i) standard nonparametric bootstrap with 100 replicates, (ii) Shimodaira–Hasegawa approximate likelihood ratio test (SH-aLRT) with 1,000 replicates, and (ii) ultrafast bootstrap with 10,000 replicates. The mitochondrial genome was retrieved from the genome assembly (see below), annotated with MitoZ (Meng et al. 2019) and compared to the mitogenome of *Gordionus alpestris* (NCBI accession number MG257765).

### Genomic DNA extraction

Entire specimens of *G. montsenyensis* were flash frozen in liquid nitrogen. High Molecular Weight (HMW) gDNA was extracted from a single specimen using Nanobind tissue kit (Circulomics), following the manufacturer’s protocol. The HMW gDNA was eluted in elution buffer, quantified by Qubit DNA BR Assay kit (Thermo Fisher Scientific) and the DNA purity was estimated from UV/Vis measurements using Nanodrop 2000 (Thermo Fisher Scientific). To determine the gDNA integrity pulse-field gel electrophoresis, using the Pippin Pulse (Sage Science) was performed. The gDNA samples were stored at 4°C until subsequent analysis.

### Long-read whole-genome library preparation and sequencing

After extraction, HMW gDNA was quality controlled for purity, quantity and integrity for long read sequencing. The sequencing libraries were prepared using the 1D Sequencing kit SQK-LSK110 from Oxford Nanopore Technologies (ONT). The sequencing run was performed on PromethIon 24 (ONT) using a flow cell R9.4. FLO-PRO002 (ONT) and the sequencing data was collected for 110 hours. The quality parameters of the sequencing runs were monitored by the MinKNOW platform version 4.3.4 in real time and basecalled with Guppy version 5.0.11.

### Short-read whole-genome sequencing library preparation and sequencing

HMW gDNA was extracted as described above from a second specimen for short-read library sequencing. The short insert paired-end libraries for the whole genome sequencing were prepared with PCR free protocol using KAPA HyperPrep kit (Roche). The library was sequenced on NovaSeq 6000 (Illumina) with a read length of 2×151 bp. Image analysis, base calling and quality scoring of the run were processed using the manufacturer’s software Real Time Analysis (RTA v3.4.4) and followed by generation of FASTQ sequence files.

### Chromatin conformation capture library preparation and sequencing

Chromatin conformation capture sequencing (Hi-C) library was prepared using the HiC High-coverage kit (Arima Genomics) following manufacturer’s instructions. Sample concentration was assessed using a Qubit DNA HS Assay kit (Thermo Fisher Scientific) and library preparation was carried out using the Accel-NGS 2S Plus DNA Library Kit (Swift Bioscience) and amplified with the KAPA HiFi DNA polymerase (Roche). The amplified library was sequenced on NovaSeq 6000 (Illumina) with a read length of 2×151 bp.

### RNA extraction, library preparation and sequencing

RNA extraction was obtained from a flash-frozen whole specimen using HigherPurity™ Total RNA Extraction Kit (Canvax Biotech) and following manufacturer’s instructions. Concentration was assessed by Qubit RNA BR Assay kit (Thermo Fisher Scientific). Samples were subjected to Illumina’s TruSeq Stranded mRNA library preparation kit and sequenced on NovaSeq 6000 (Illumina, 2×151 bp). In total ∼127 million reads were generated.

### Preprocessing

The genome size and the heterozygosity of the *Gordionus montsenyensis* sp. nov. was estimated based on *k*-mer distribution of the short DNA Illumina reads using Jellyfish v2.3.0 (Marçais and Kingsford 2011), along with GenomeScope v2.0 (Ranallo-Benavidez, Jaron, and Schatz 2020) with *k*=27 (Figure 4).

Quality assessment of the raw DNA Illumina reads was performed with FastQC v0.11.9 (https://www.bioinformatics.babraham.ac.uk/projects/fastqc/). The adapters and reads less than 10 bp (option -m 10) were trimmed using cutadapt v2.9 (Martin 2011). Quality assessment of the raw Nanopore reads was performed by NanoPack v1.39.0 (De Coster et al. 2018). Adapter trimming and length filtering of Nanopore reads was done using Porechop v0.2.4 (https://github.com/rrwick/Porechop) with --min_split_read_size 5. The filtered Illumina reads were used as reference in Filtlong v0.2.0 (https://github.com/rrwick/Filtlong) to filter the trimmed Nanopore data and the chimeric reads were removed using yacrd v1.0.0 (Marijon, Chikhi, and Varré 2020). Ratatosk v0.7.5 (Holley et al. 2021) was employed for hybrid error correction of the Nanopore reads as the last step of the preprocessing. The pipeline followed for the trimming of Hi-C reads was the same as the short Illumina reads.

RNA-seq reads were assessed with FastQC and the low-quality reads and adapters were removed using Trimmomatic v0.39 (Bolger, Lohse, and Usadel 2014) (MINLEN: 75, SLIDINGWINDOW: 4:15, LEADING: 10, TRAILING: 10, AVGQUAL: 30).

All the FastQC output file of the raw and preprocessed data of Illumina, Hi-C and RNA-seq can be found in https://github.com/MetazoaPhylogenomicsLab/ChromosomeLevelGenomes/blob/main/Gordionus_montsenyensis/Gordionus_montsenyensis_QC.tar.gz.

### Genome assembly

The initial de novo genome assembly was generated with the previously described preprocessed Nanopore reads using the Flye v2.9 (Kolmogorov et al. 2019) assembler with default parameters. To remove superfluous haplotigs, we purged the initial assembly using minimap2 v2.24 (Li 2018) with parameter -x map-ont and purge_dups v1.0.1 (Guan et al. 2020). The obtained sequences were further scaffolded into a chromosome-level assembly by integrating the Hi-C sequencing data using instaGRAAL v0.1.6 no-opengl branch (Baudry et al. 2020) with parameters -l 4 -n 50 and the contact map was built by aligning the Hi-C reads to the genome assembly with bowtie2 v2.3.5.1 (Langmead et al. 2019) and the hicstuff pipeline v3.1.4 (Matthey-Doret et al. 2020) with parameters -e DpnII,HinfI -m iterative. To fill in the gaps, TGS-GapCloser v1.1.1 (Xu et al. 2020) was used with parameters --ne --tgstype ont. Manual curation was performed using the GRIT Rapid Curation workflow (https://gitlab.com/wtsi-grit/rapid-curation) (Howe et al. 2021), Pretext (https://github.com/wtsi-hpag/PretextView) and the visualization of the contact maps with the hicstuff pipeline.

The mitochondrial genome was assembled with MITGARD (Nachtigall, Grazziotin, and Junqueira-de-Azevedo 2021) using the WGS Illumina paired-end reads and the mitochondria of *Gordionus alpestris* (NC_044095.1) as a reference and selecting the clade Arthropoda. The mitochondrial genome was annotated using MitoZ with parameters annotate --genetic_code auto --clade Arthropoda (Meng et al. 2019). The mitochondrial genome was aligned using NCBI Blast with the megablast algorithm against the non-redundant nucleotide database (nt) with default parameters.

All the resulting assembly statistics were calculated with assembly-stats (https://github.com/rjchallis/assembly-stats) and assessed with BUSCO v5.4.7 (Manni et al. 2021) against the metazoa_odb10 reference set in genome mode. *k*-mer analysis of the final assembly was done using KAT v2.4.2 (Mapleson et al. 2017) and the module kat comp against the Illumina reads. Haploidy score was calculated by mapping the Nanopore reads to the assembly with minimap2 v2.24 and using HapPy v0.1 (Guiglielmoni et al. 2021) with parameter -s 280M.

### Repeat identification

Repetitive elements of the genome of *Gordionus montsenyensis* sp. nov. were identified by building a de novo library using RepeatModeler2 v2.0.3 (-e rmblast) (Flynn et al. 2020), along with the extra LTRStruct pipeline. The software RepeatMasker v4.1.2-p1 (-a -s -xsmall -e rmblast -pa 5 -nolow -norna -no_is -gff) (Tarailo-Graovac and Chen 2009) was used to annotate and soft mask the repetitive elements, based on the previously generated de novo library and the incorporated, in RepeatMasker, Dfam curated v3.6 database.

### Gene prediction

After masking the repetitive elements, protein coding gene annotation was performed by integrating ab-initio and homology-based evidence with BRAKER v2.1.6. First, the preprocessed RNA-seq data were aligned to the manually curated genome assembly of *Gordionus montsenyensis* sp. nov. using STAR v2.7.9a (Dobin et al. 2013) and the aligned BAM file was used as input in BRAKER1 (Hoff et al. 2016), which incorporates GeneMark-ET (Lomsadze, Burns, and Borodovsky 2014) together with AUGUSTUS v3.4.0 (Stanke et al. 2008) to train and predict gene structures. All metazoan proteins of the OrthoDB v10 along with protein sequences from *Gordius* sp. KK-2019 (SRR8943410), *Paragordius varius* (ERR1817118)*, Nectonema munidae* (SRR17843037) were aligned against the manually curated genome assembly using the ProtHint pipeline v2.6.0 (Brůna, Lomsadze, and Borodovsky 2020) and incorporated in the BRAKER2 (Brůna et al. 2021) with GeneMark-EP+ (Brůna, Lomsadze, and Borodovsky 2020) and AUGUSTUS, as evidence for homology-based prediction. Protein sequences from *Gordius* sp. KK-2019, *Paragordius varius* and *Nectonema munidae* were obtained and de novo assembled as described in MATEdb (Fernández et al. 2022). Both BRAKER1 and BRAKER2 annotations were combined and filtered with TSEBRA (Gabriel et al. 2021), generating the final set of predicted protein-coding genes.

## Data Availability

The raw genomic and transcriptomic data along with the genome assembly sequence and annotation of *Gordionus montsenyensis* sp. nov. are deposited in the European Nucleotide Archive, the later also available in the GitHub repository from the Metazoa Phylogenomics Lab (https://github.com/MetazoaPhylogenomicsLab/ChromosomeLevelGenomes/upload/main/Gordionus_montsenyensis) and their corresponding accession numbers are available in Table 2.

## Author Contribution

KE, JSO and RF designed and conceived the research. GIMR and RF collected the samples. JSO extracted nucleic acids and constructed the Hi-C library. MG supervised the sequencing of all libraries at the Centro Nacional de Análisis Genómico (CNAG, Barcelona). KE and NG assembled the genome. KE and GIMR annotated the genome. CV assembled and annotated the mitogenome. JFF provided advice on genome assembly. ASR and RF formally described the species. RF supervised the study. KE and RF drafted the paper. All authors contributed to the final version of the manuscript.

## Acknowledgements

We thank the members of the Metazoa Phylogenomics Lab for their support during the collection of the specimens (Vanina Tonzo, Jesús Lozano Fernández, Pau Balart and Leandro Aristide). We are thankful to Jonathan Wood and his team for the help with the second round of manual curation of the assembly. We would like to acknowledge and thank all supplier partners that have kindly donated kits, reagents to the ERGA pilot Library Preparation Hubs to support species to produce the generation of high-quality genomes and annotations. This support has been key to embedding a culture of diversity, equity, inclusion, and justice in the Pilot Project. Specifically we want to thank Arima Genomics, Integrated DNA Technologies (IDT), Agilent Technologies, Fisher Scientific Spain and Illumina Inc. We acknowledge access to the storage resources at Barcelona Supercomputing Center, which are partially funded from European Union H2020-INFRAEOSC-2018-2020 programme through the DICE project (Grant Agreement no. 101017207). RF acknowledges support from the following sources of funding: Ramón y Cajal fellowship (grant agreement no. RYC2017-22492 funded by MCIN/AEI/10.13039/501100011033 and ESF ‘Investing in your future’), the Agencia Estatal de Investigación (project PID2019-108824GA-I00 funded by MCIN/AEI/10.13039/501100011033) and the European Research Council (this project has received funding from the European Research Council (ERC) under the European’s Union’s Horizon 2020 research and innovation programme (grant agreement no. 948281). We also thank Centro de Supercomputación de Galicia (CESGA) for access to computer resources. This genome is part of the ERGA Pilot project from Spain. We thank the ERGA pilot executive committee (Ann Mc Cartney, Alice Mouton and Giulio Formenti) for the coordination of the ERGA pilot project and the continuous support during the establishment phase of ERGA.

## Notes

### Competing Interest Statement

The authors have declared no competing interest.

### Summary of Updates

Version recommended by PCI Genomics. Added the badge as required to show it.

## References

Aguinaldo, Anna Marie A., James M. Turbeville, Lawrence S. Linford, Maria C. Rivera, J. R. Garey, R. A. Raff, and J. A. Lake. 1997. “Evidence for a Clade of Nematodes, Arthropods and Other Moulting Animals.” Nature 387, no. 6632 (May 1, 1997): 489–93. 10.1038/387489a0.

Baudry, Lyam, Nadège Guiglielmoni, Hervé Marie-Nelly, Alexandre Cormier, Martial Marbouty, Komlan Avia, Mie Yl, et al. “instaGRAAL: Chromosome-Level Quality Scaffolding of Genomes Using a Proximity Ligation-Based Scaffolder.” Genome Biology 21, no. 1 (June 18, 2020). 10.1186/s13059-020-02041-z.

Begay, Alyssa C., Andreas Schmidt-Rhaesa, Matthew G. Bolek, and Ben Hanelt. “Two New *Gordionus* Species (Nematomorpha: Gordiida) from the Southern Rocky Mountains (USA).” Zootaxa 3406, no. 1 (August 1, 2012): 30. 10.11646/zootaxa.3406.1.2.

Blaxter, Mark, Paul De Ley, James R. Garey, Leo X. Liu, Patsy Scheldeman, Andy Vierstraete, Jacques R. Vanfleteren, et al. “A Molecular Evolutionary Framework for the Phylum Nematoda.” Nature 392, no. 6671 (March 1, 1998): 71–75. 10.1038/32160.

Bolger, Anthony M., Marc Lohse, and Björn Usadel. “Trimmomatic: A Flexible Trimmer for Illumina Sequence Data.” Bioinformatics 30, no. 15 (April 1, 2014): 2114–20. 10.1093/bioinformatics/btu170.

Brůna, Tomáš, Katharina J. Hoff, Alexandre Lomsadze, Mario Stanke, and Mark Borodovsky. “BRAKER2: Automatic Eukaryotic Genome Annotation with GeneMark-EP+ and AUGUSTUS Supported by a Protein Database.” NAR Genomics and Bioinformatics 3, no. 1 (January 6, 2021). 10.1093/nargab/lqaa108.

Brůna, Tomáš, Alexandre Lomsadze, and Mark Borodovsky. “GeneMark-EP+: Eukaryotic Gene Prediction with Self-Training in the Space of Genes and Proteins.” NAR Genomics and Bioinformatics 2, no. 2 (May 13, 2020). 10.1093/nargab/lqaa026.

Cunha, T. J., de Medeiros, B. A. S., Lord, A., Sørensen, M. V., & Giribet, G. (2023). Rampant loss of universal metazoan genes revealed by a chromosome-level genome assembly of the parasitic Nematomorpha. Current Biology: CB, 33(16), 3514–3521.e4. 10.1016/j.cub.2023.07.003

De Coster, Wouter, Svenn D’Hert, Darrin T. Schultz, Marc Cruts, and Christine Van Broeckhoven. “NanoPack: Visualizing and Processing Long-Read Sequencing Data.” Bioinformatics 34, no. 15 (March 14, 2018): 2666–69. 10.1093/bioinformatics/bty149.

De Villalobos, L. C., Ignacio Ribera, and Ian Downie. “Hairworms Found in Scottish Agricultural Land, with Descriptions of Two New Species of *Gordionus Muller* (Nematomorpha: Gordiidae).” Journal of Natural History 33, no. 12 (December 1, 1999): 1767–80. 10.1080/002229399299716.

Dobin, Alexander, Carrie A. Davis, Felix Schlesinger, Jorg Drenkow, Chris Zaleski, Sonali Jha, Philippe Batut, Mark Chaisson, and Thomas R. Gingeras. “STAR: Ultrafast Universal RNA-Seq Aligner.” Bioinformatics 29, no. 1 (October 25, 2012): 15–21. 10.1093/bioinformatics/bts635.

Fernández, Rosa, Vanina Tonzo, Carolina Márquez Guerrero, Jesus Lozano-Fernandez, Gemma I. Martínez-Redondo, Pau Balart-García, Leandro Aristide, Klara Eleftheriadi, and Carlos Vargas-Chávez. “MATEdb, a Data Repository of High-Quality Metazoan Transcriptome Assemblies to Accelerate Phylogenomic Studies.” Peer Community Journal 2 (October 5, 2022). 10.24072/pcjournal.177.

Flynn, Jullien M., Robert Hubley, Clément Goubert, Jeb Rosen, Andrew G. Clark, Cédric Feschotte, and Arian F.A. Smit. “RepeatModeler2 for Automated Genomic Discovery of Transposable Element Families.” Proceedings of the National Academy of Sciences of the United States of America 117, no. 17 (April 16, 2020): 9451–57. 10.1073/pnas.1921046117.

Folmer, O. F., Michael J. Black, Walter R. Hoeh, Richard A. Lutz, and Robert C. Vrijenhoek. “DNA Primers for Amplification of Mitochondrial Cytochrome c Oxidase Subunit I from Diverse Metazoan Invertebrates.” PubMed 3, no. 5 (October 1, 1994): 294–99. https://pubmed.ncbi.nlm.nih.gov/7881515.

Gabriel, Lars, Katharina J. Hoff, Tomáš Brůna, Mark Borodovsky, and Mario Stanke. “TSEBRA: Transcript Selector for BRAKER.” BMC Bioinformatics 22, no. 1 (November 25, 2021). 10.1186/s12859-021-04482-0.

Gerlach, S.A. (1978): Nematomorpha. In: Limnofauna Europaea. Illies, J. (ed.). Gustav Fischer Verlag, Stuttgart: 50–53.

Guan, Dengfeng, Shane A. McCarthy, Jonathan Wood, Kerstin Howe, Yadong Wang, and Richard Durbin. “Identifying and Removing Haplotypic Duplication in Primary Genome Assemblies.” Bioinformatics 36, no. 9 (January 23, 2020): 2896–98. 10.1093/bioinformatics/btaa025.

Guiglielmoni, Nadège, Antoine Houtain, Alessandro Derzelle, Karine Van Doninck, and Jean-François Flot. “Overcoming Uncollapsed Haplotypes in Long-read Assemblies of Non-Model Organisms.” BMC Bioinformatics 22, no. 1 (June 5, 2021). 10.1186/s12859-021-04118-3.

Hanelt, Ben, Francis T. Thomas, and Andreas Schmidt-Rhaesa. “Biology of the Phylum Nematomorpha.” Advances in Parasitology Volume 59,243–305, 2005. 10.1016/s0065-308x(05)59004-3.

Hanelt, Ben, and John J. Janovy. “Spanning the Gap: Experimental Determination of Paratenic Host Specificity of Horsehair Worms (Nematomorpha: Gordiida).” Invertebrate Biology 122, no. 1 (May 11, 2005): 12–18. 10.1111/j.1744-7410.2003.tb00068.x.

Hoff, Katharina J., Simone Lange, Alexandre Lomsadze, Mark Borodovsky, and Mario Stanke. “BRAKER1: Unsupervised RNA-seq-Based Genome Annotation with GeneMark-ET and AUGUSTUS.” Bioinformatics 32, no. 5 (November 11, 2015): 767–69. 10.1093/bioinformatics/btv661.

Holley, Guillaume, Doruk Beyter, Helga Ingimundardottir, Peter Møller, Snaedis Kristmundsdottir, Hannes P. Eggertsson, and Bjarni V. Halldorsson. “Ratatosk: Hybrid Error Correction of Long Reads Enables Accurate Variant Calling and Assembly.” Genome Biology 22, no. 1 (January 8, 2021). 10.1186/s13059-020-02244-4.

Howe, Kerstin, William Chow, Joanna Collins, Sarah Pelan, Damon-Lee Pointon, Ying Sims, James Torrance, Alan S. Tracey, and Jonathan Wood. “Significantly Improving the Quality of Genome Assemblies through Curation.” GigaScience 10, no. 1 (January 1, 2021). 10.1093/gigascience/giaa153.

Kalyaanamoorthy, Subha, Bui Quang Minh, Thomas K.S. Wong, Arndt Von Haeseler, and Lars S. Jermiin. “ModelFinder: Fast Model Selection for Accurate Phylogenetic Estimates.” Nature Methods 14, no. 6 (May 8, 2017): 587–89. 10.1038/nmeth.4285.

Katoh, Kazutaka, and Daron M. Standley. 2013. “MAFFT Multiple Sequence Alignment Software Version 7: Improvements in Performance and Usability.” Molecular Biology and Evolution 30 (4): 772–80.

Kolmogorov, Mikhail, Jeffrey Yuan, Yu Lin, and Pavel A. Pevzner. “Assembly of Long, Error-Prone Reads Using Repeat Graphs.” Nature Biotechnology 37, no. 5 (April 1, 2019): 540–46. 10.1038/s41587-019-0072-8.

Langmead, Ben, Christopher Wilks, Valentin Antonescu, and Rone Charles.“Scaling Read Aligners to Hundreds of Threads on General-Purpose Processors.” Bioinformatics 35, no. 3 (July 18, 2018): 421–32. 10.1093/bioinformatics/bty648.

Laumer, Christopher E., Rosa Fernández, Sarah Lemer, David Combosch, Kevin M. Kocot, Ana Riesgo, Sónia C. S. Andrade, Wolfgang Sterrer, Martin V. Sørensen, and Gonzalo Giribet. “Revisiting Metazoan Phylogeny with Genomic Sampling of All Phyla.” Proceedings of the Royal Society B: Biological Sciences 286, no. 1906 (July 10, 2019): 20190831. 10.1098/rspb.2019.0831.

Li, Heng. “Minimap2: Pairwise Alignment for Nucleotide Sequences.” Bioinformatics 34, no. 18 (May 10, 2018): 3094–3100. 10.1093/bioinformatics/bty191.

Lomsadze, Alexandre, Paul Burns, and Mark Borodovsky. “Integration of Mapped RNA-Seq Reads into Automatic Training of Eukaryotic Gene Finding Algorithms.” Nucleic Acids Research 42, no. 15 (July 2, 2014): e119. 10.1093/nar/gku557.

Manni, Mosè, Matthew R Berkeley, Mathieu Seppey, and Evgeny M. Zdobnov. “BUSCO: Assessing Genomic Data Quality and Beyond.” Current Protocols 1, no. 12 (December 1, 2021). 10.1002/cpz1.323.

Mapleson, Daniel, Gonzalo Garcia Accinelli, George Kettleborough, Jonathan Wright, and Bernardo J. Clavijo. “KAT: A K-Mer Analysis Toolkit to Quality Control NGS Datasets and Genome Assemblies.” Bioinformatics 33, no. 4 (November 28, 2016): 574–76. 10.1093/bioinformatics/btw663.

Marçais, Guillaume, and Carl Kingsford. “A Fast, Lock-Free Approach for Efficient Parallel Counting of Occurrences of *k*-Mers.” Bioinformatics 27, no. 6 (January 7, 2011): 764–70. 10.1093/bioinformatics/btr011.

Marijon, Pierre, Rayan Chikhi, and Jean-Stéphane Varré. “Yacrd and Fpa: Upstream Tools for Long-Read Genome Assembly.” Bioinformatics 36, no. 12 (April 21, 2020): 3894–96. 10.1093/bioinformatics/btaa262.

Martin, Marcel. “Cutadapt Removes Adapter Sequences from High-Throughput Sequencing Reads.” EMBnet Journal 17, no. 1 (May 2, 2011): 10. 10.14806/ej.17.1.200.

Matthey-Doret, Cyril, Lyam Baudry, Amaury Bignaud, Rémi Montagne, Nadège Guiglielmoni, Théo Foutel-Rodier, and Vittore F. Scolari. “hicstuff: Simple library/pipeline to generate and handle Hi-C data.” Zenodo, 2020. 10.5281/zenodo.4066363.

Minh, Bui Quang, Heiko Schmidt, Olga Chernomor, Dominik Schrempf, Michael D. Woodhams, Arndt Von Haeseler, and Robert Lanfear. “IQ-TREE 2: New Models and Efficient Methods for Phylogenetic Inference in the Genomic Era.” Molecular Biology and Evolution 37, no. 5 (February 3, 2020): 1530–34. 10.1093/molbev/msaa015.

Meng, Guanliang, Yiyuan Li, Chentao Yang, and Shanlin Liu. “MitoZ: A Toolkit for Animal Mitochondrial Genome Assembly, Annotation and Visualization.” Nucleic Acids Research 47, no. 11 (March 13, 2019): e63. 10.1093/nar/gkz173.

Nachtigall, Pedro G., Felipe G. Grazziotin, and Inácio L.M. Junqueira-De-Azevedo. “MITGARD: An Automated Pipeline for Mitochondrial Genome Assembly in Eukaryotic Species Using RNA-Seq Data.” Briefings in Bioinformatics 22, no. 5 (January 30, 2021). 10.1093/bib/bbaa429.

Ranallo-Benavidez, T. Rhyker, Kamil S. Jaron, and Michael C. Schatz. “GenomeScope 2.0 and Smudgeplot for Reference-Free Profiling of Polyploid Genomes.” Nature Communications 11, no. 1 (March 18, 2020). 10.1038/s41467-020-14998-3.

Schmidt-Rhaesa, A., and A. Cieslak. “Three new species of Paragordionus and Gordionus (Nematomorpha, Gordiida) from Spain and Turkey, with comments on the taxon Semigordionus.” Mitteilungen aus dem Hamburgischen Zoologischen Museum und Institut 105 (2008): 13–22.

Schmidt-Rhaesa, Andreas, and Jesús Martínez. “*Gordius gonzalezi*, a new Species of Horsehair Worms (Nematomorpha) from Spain.” Zootaxa, April 11, 2016. 10.11646/zootaxa.4103.1.6.

Schmidt-Rhaesa, Andreas, and Valeria Viktoria Gusich. “A new *Gordionus* MÜLLER, 1927 from Switzerland (Nematomorpha, Gordiida).” Revue Suisse De Zoologie 117 (January 1, 2010): 77–82. 10.5962/bhl.part.117589.

Schmidt-Rhaesa, Andreas, Ben Hanelt, and Will K. Reeves. “Redescription and Compilation of Nearctic Freshwater Nematomorpha (Gordiida), with the Description of Two New Species.” Proceedings of the Academy of Natural Sciences of Philadelphia 153, no. 1 (December 1, 2003): 77–117. 10.1635/0097-3157(2003)153.

Schmidt-Rhaesa, Andreas, Monica A. Farfan, and Ernest C. Bernard. “First Record of Millipeds as Hosts for Horsehair Worms (Nematomorpha) in North America.” Northeastern Naturalist 16, no. 1 (March 1, 2009): 125–30. 10.1656/045.016.0110.

Schmidt-Rhaesa, Andreas. Nematomorpha, Priapulida, Kinorhyncha, Loricifera. Walter de Gruyter, 2013.

Smith, Douglas H. “Observations on the Morphology and Taxonomy of Two *Parachordodes* species (Nematomorpha, Gordioida, Chordodidae) in Southern New England (USA).” Journal of Zoology 225, no. 3 (November 1, 1991): 469–80. 10.1111/j.1469-7998.1991.tb03829.x.

Stanke, Mario, Mark Diekhans, Robert Baertsch, and David Haussler. “Using Native and Syntenically Mapped cDNA Alignments to Improve de Novo Gene Finding.” Bioinformatics 24, no. 5 (January 24, 2008): 637–44. 10.1093/bioinformatics/btn013.

Sørensen, Martin V., Martin B. Hebsgaard, Iben Heiner, Henrik Glenner, Eske Willerslev, and Reinhardt Møbjerg Kristensen. “New Data from an Enigmatic Phylum: Evidence from Molecular Sequence Data Supports a Sister-Group Relationship between Loricifera and Nematomorpha.” Journal of Zoological Systematics and Evolutionary Research 46, no. 3 (August 1, 2008): 231–39. 10.1111/j.1439-0469.2008.00478.x.

Tarailo-Graovac, Maja, and Nansheng Chen. “Using RepeatMasker to Identify Repetitive Elements in Genomic Sequences.” Current Protocols in Bioinformatics 25, no. 1 (March 1, 2009). 10.1002/0471250953.bi0410s25.

Thomas, Francis T., Andreas Schmidt-Rhaesa, Guilhaume Martin, C. Manu, Patrick Durand, and Florent Renaud. “Do Hairworms (Nematomorpha) Manipulate the Water Seeking Behavior of Their Terrestrial Hosts?” Journal of Evolutionary Biology 15, no. 3 (April 30, 2002): 356–61. 10.1046/j.1420-9101.2002.00410.x.

Xu, Mengyang, Lidong Guo, Shengqiang Gu, Ou Wang, Rui Zhang, Brock A. Peters, Guangyi Fan, et al. “TGS-GapCloser: A Fast and Accurate Gap Closer for Large Genomes with Low Coverage of Error-Prone Long Reads.” GigaScience 9, no. 9 (September 1, 2020). 10.1093/gigascience/giaa094.

